# More than a token photo: humanizing scientists enhances student engagement

**DOI:** 10.1101/2024.01.29.577791

**Authors:** Robin A. Costello, Emily P. Driessen, Melissa K. Kjelvik, Elizabeth H. Schultheis, Rachel M. Youngblood, Ash T. Zemenick, Marjorie G. Weber, Cissy J. Ballen

## Abstract

Highlighting scientists from historically excluded groups in educational materials increases student engagement in STEM. However, which specific elements of these educational materials maximize their impact remains untested, leaving educators guessing how to best highlight counter-stereotypical scientists in their classrooms. We tested the effects of including visual and humanizing descriptions of scientists featured in quantitative biology activities on over 3,700 students across 36 undergraduate institutions. We found that including humanizing information about counter-stereotypical scientists increased the extent to which students related to those scientists, which in turn translated to higher student engagement. Students who shared one or more excluded identity(s) with the featured scientists related most strongly. Our findings demonstrate the importance of humanizing counter-stereotypical scientists in classrooms, beyond simply adding a photo to increase representation.

## Introduction

Undergraduate students in science, technology, engineering, and mathematics (STEM) are unlikely to see a diversity of scientists represented in their educational materials (*1–5*). The mismatch between the identities of scientists featured in courses and an increasingly diversified student body (*6*) upholds the stereotype of scientists as white, cis-men, perpetuating the false narrative that only certain types of people contribute to the scientific enterprise (*7–10*).

Educators and curriculum developers have worked to combat this false narrative by creating educational materials that highlight scientists from a diversity of backgrounds, intentionally featuring scientists whose identities do not match the dominant stereotype of a scientist (hereafter, counter-stereotypical scientists) (*11–15*). Featuring counter-stereotypical scientists in course materials is billed as a scalable, accessible, and easy-to-implement tool that increases the recruitment and retention of students with identities historically and currently excluded from STEM (*11*). However, although early evidence has demonstrated the efficacy of classroom interventions that feature counter-stereotypical scientists (*12–17*), the specific elements of these interventions that are most impactful remain unclear (*18, 19*). The lack of research in this area leaves educators and curriculum developers without data-supported guidelines for how to effectively develop and employ classroom materials that feature counter-stereotypical scientists.

Theory predicts that counter-stereotypical scientists must be *relatable* for students to engage with STEM content and pursue STEM careers (*20*). Students are hypothesized to relate to scientists along two different axes: shared visual elements (i.e., that scientist looks like me) and shared humanizing elements (i.e., that scientist shares similar values, hobbies, and experiences with me) (*20*). Here, we test this theory by asking whether a photo is enough to make featured scientists relatable or whether humanizing elements are necessary. Specifically, we employed a large-scale experimental approach in undergraduate biology courses across the United States to parse the effects of visual and humanizing elements of classroom materials featuring counter-stereotypical scientists on how students relate to scientists and engage with STEM.

In a nationwide, multi-institutional study, we experimentally manipulated the way biology activities highlight counter-stereotypical scientists. Students were given one of three versions of short quantitative exercises (Data Nuggets, *21, 22*) that feature data from contemporary scientists who self-report as having identities excluded from STEM (Project Biodiversify, *23*). As treatments, students were exposed to either: quantitative exercises without any information about the scientist (control), b) quantitative exercises with photos of the scientist (visual treatment), and c) quantitative exercises with scientist photos and interview questions answered by the scientist about their experiences as counter-stereotypical scientists (humanizing treatment) (Fig. 1A). Treatments were implemented by 43 instructors across 36 U.S. undergraduate institutions, representing 3,788 students (SI Appendix, Fig. S1). At the conclusion of each of the three activities, students responded to survey items measuring the extent to which they related to the featured scientist and their engagement in the quantitative biology activity (*24, 25*).

**Fig. 1.**
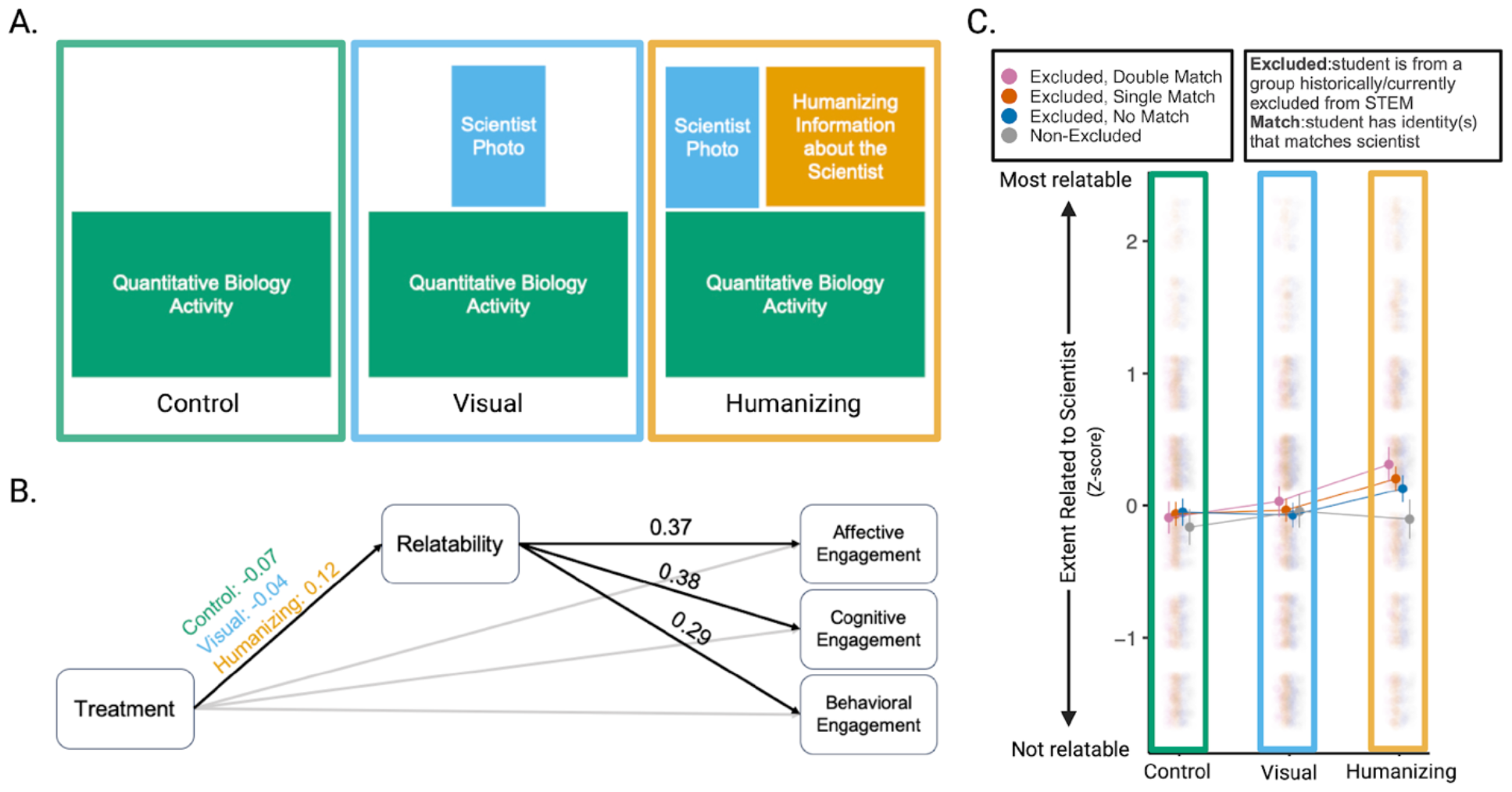
Effects of visual and humanizing elements on how students relate to scientists and engage with quantitative biology activities. (**A**) Three treatments used in this study varied the extent to which visual and humanizing elements were provided during quantitative biology activities. (**B**) Our path model found that the perceived relatability of the scientist mediated the relationship between the three treatments and all three dimensions of student engagement on the quantitative biology activities. In particular, the humanizing treatment led to higher scientist relatability which increased student engagement. Black paths are significant (P < 0.05). Marginal means are reported for the path connecting treatment to scientist relatability, and mean-variance standardized beta coefficients are reported for paths connecting scientist relatability to the three components of engagement. (**C**) Marginal means with 95% confidence error bars of the extent to which students related to the featured scientist by treatment and the extent to which the student gender and race/ethnicity identities matched the identities of the scientist featured. Jittered points (N = 6,244 survey responses from 3,102 students) depict the extent to which students related to the scientist on a 7-point sliding scale, ranging from 1 (not at all) to 7 (very), that has been mean and variance standardized.

## Results and Discussion

### What Enhances Student Engagement?

Path analyses showed that when quantitative biology activities included humanizing information about counter-stereotypical scientists (humanizing treatment), students related more to the scientists (P < 0.0001; N = 7,361 survey responses from 3,788 students; Fig. 1B; SI Appendix, Table S1). Further, relating more to humanized scientists was associated with higher student engagement with the quantitative biology activities. This was true across three proxy measures for student engagement: a) interest in the activities (affective engagement; *β* = 0.37; P < 0.0001), b) perceived importance of the activities (cognitive engagement; *β* = 0.38; P < 0.0001), and c) effort spent on the activities (behavioral engagement; *β* = 0.29; P < 0.0001; Fig. 1B; SI Appendix, Table S1). These results illustrate that humanizing information about counter-stereotypical scientists increases the extent to which students relate to those scientists, which in turn translates to higher student engagement. Simply describing research or including photos of scientists is not enough to increase scientist relatability and student engagement.

### Does it Matter Whether Students Share the Same Identities as Scientists?

Exposure to curricular materials that feature counter-stereotypical scientists is expected to shift attitudes towards science and scientists, especially among students who share identities with the featured scientist (*18, 19*). To test this hypothesis, we demographically matched students and scientists based on whether they shared gender and/or race/ethnicity identities, as indicated by survey responses. Students who shared one or more excluded identity(s) with the featured scientist rated those scientists as most relatable when activities included humanizing information (*double match* vs. *non-excluded*: Cohen’s *d* = 0.331, P = 0.0005; *single match* vs. *non-excluded*: Cohen’s *d* = 0.245, P = 0.007; N = 6,244 survey responses from 3,102 students; Fig. 1C; SI Appendix, Table S2). Conversely, activities that included only visual elements of the featured scientists were not sufficient to create change in how students with shared excluded identity(s) related to the scientists (*double match* vs. *non-excluded*: Cohen’s *d* = 0.060, P = 0.999; *single match* vs. *non-excluded*: Cohen’s *d* = 0.006, P = 1.00; Fig. 1C; SI Appendix, Table S2). The control treatment likewise found no relationship between shared excluded identity(s) and scientist relatability (*double match* vs. *non-excluded*: Cohen’s *d* = 0.057, P = 1.00; *single match* vs. *non-excluded*: Cohen’s *d* = 0.079, P = 0.969; Fig. 1C; SI Appendix, Table S2). Our results emphasize that including humanizing information about counter-stereotypical scientists in course materials is necessary for students to relate to scientists with whom they share excluded identities.

### What do Students Relate to?

To better understand what humanizing features students relate to, we asked students in the humanizing treatment to describe *how* they related to the featured scientists. Qualitative data analysis found that 20.5% of students related to the excluded identity(s) held by the featured scientists, including the scientists’ race/ethnicity, gender, age, and sexuality (SI Appendix, Fig. S2; SI Appendix, Fig. S3). Results further supported the importance of identity sharing: students with shared excluded identity(s) had over 28 times higher odds of reporting relating to the excluded identities of the featured scientists compared to white men students (*double match*: odds ratio = 38.981; P < 0.0001; *single match*: odds ratio = 28.807; P < 0.0001; N = 1,324 responses from 742 students; SI Appendix, Fig. S3; SI Appendix, Table S3). Students with excluded identity(s) but not identity(s) that specifically matched those of the featured scientist still had 5 times higher odds than white men students of mentioning relating to the excluded identities of the featured scientist (*no match*: odds ratio = 5.354, P = 0.0013; SI Appendix, Fig. S3; SI Appendix, Table S3). This qualitative analysis reiterates the importance of ‘seeing yourself in science’, as students related most to the identities of the scientist when they shared the same excluded identities as scientists. However, our results also reveal that students still relate to the counter-stereotypical identities held by scientists, even when those students do not share the same excluded identities as the featured scientist.

### In Conclusion: Embrace the Humanity of Scientists

Three key findings emerge from our research. 1) Including humanizing information about the values, hobbies, and experiences of counter-stereotypical scientists featured in curricular materials is necessary for students to relate to scientists. 2) Students who share one or more excluded identity(s) with humanized scientists relate most strongly to featured scientists. 3) When students relate to scientists featured in curricular materials, they engage more with the materials.

Our results demonstrate that including photos of scientists in course materials is not sufficient to change student attitudes about science. Rather, to engage students, and especially those with excluded identities, it is critical we also humanize scientists. Instructors can enact this change by implementing existing humanizing resources in their courses, such as DataVersify (*21, 22*), Project Biodiversify (*23*), Scientist Spotlights (*8–17*), BioGraphI, Story Collider Podcast, and SACNAS Biography Project. These curricular materials all vary widely in how they incorporate humanizing information, differing in the amount of information included and the way that information is conveyed (i.e., through text, audio, video). Identifying the types of humanizing information commonly provided to students and isolating their relative impact is an important next step for curricular resource development.

Our findings show that it is vital to intentionally center and humanize counter-stereotypical scientists in curricular materials. By embracing the humanity of scientists, curricular materials can signal to students that they too can contribute to STEM fields. Our work provides a simple approach that curriculum developers and instructional/departmental policies can employ to improve equity in STEM.

## Materials and Methods

### Experimental Design

We used a multi-level design in which instructors of undergraduate biology courses for majors and non-majors in the United States applied three randomized treatments across academic terms (Fig. 1A). The three treatments were designed to assess the impacts of pairing quantitative biology activities that varied in the extent to which they shared information about featured counter-stereotypical scientists. Each academic term included the implementation of three activities from a single treatment, which were embedded within typical biology instruction and emphasized interpreting data collected by the featured scientists. Our materials were written based on the science, data, and lived experiences of 12 scientists that self-report possessing excluded backgrounds and identities (SI Appendix, Table S4) and were implemented in undergraduate institutions across 20 states (SI Appendix, Table S5). The SI Appendix contains more details on how we developed the curricular materials, manipulated scientist profiles to create our three treatments, recruited instructors, and worked with instructors to match the activities to their course content.

### Student Outcome Measures

We surveyed students with short, online Qualtrics surveys to collect demographic information (SI Appendix, Table S6) and to measure how students related to the featured scientists and engaged with the activities (SI Appendix, Table S7). To quantify scientist relatability and student engagement (including subscales for affective, cognitive, and behavioral engagement), we developed and adapted survey items that used the Experience Sampling Method, a survey approach that asks students to report their thoughts and feelings immediately following exposure to the activity (SI Appendix, Table S7) (*24, 25*). All survey items asked students to respond to the survey questions on a 7-point sliding scale, ranging from 1 (not at all) to 7 (very). To understand *how* students related to scientists, we asked students to respond to the open-ended prompt: “Describe how you related to the featured scientist in the activity, if at all”. We measured the validity of our engagement construct using confirmatory factor analysis (CFA) (SI Appendix) (*26*).

### Data Analyses

We used path models and linear mixed models to explore the effects of visual and humanizing elements of featured scientists on how students related to scientists and engaged with the activities. In all of our models, we mean-variance standardized our measures of scientist relatability and student engagement (SI Appendix). Furthermore, as classroom contexts have the potential to modulate the impacts of curricular interventions, all of our models included both the course and the specific activity implemented as random effects (*27*). We also included student ID nested within the course to account for repeated measures and avoid pseudoreplication.

First, we employed piecewise SEM of path models to examine whether student engagement with the activities was mediated by how relatable students found the featured scientists across different levels of exposure to the featured scientist (i.e., control, visualizing, or humanizing treatments) (*28*). We evaluated two separate models for each component of engagement: the full mediation model tested whether scientist relatability fully mediated the relationship between treatment and student engagement, whereas the partial mediation model additionally incorporated a direct effect between treatment and engagement (SI Appendix, Fig. S4). We fit our path analyses with the R package *piecewiseSEM* (*29*). Our path analyses included 7,361 student responses from 3,788 students (SI Appendix).

Next, we used linear mixed models to test how sharing counter-stereotypical identities with featured scientists impacts the extent to which students relate to scientists across the three treatments. We created four categories of student-scientist demographic matching with a focus on excluded identities, which we define as non-cis-man genders and non-white races (SI Appendix). If the student did *not* possess an excluded identity (i.e., was a white cis-gender man), that student was included in the Non-Excluded category. If the student held excluded gender and/or race/ethnicity identity(s), that student either shared both identities (Excluded, Double Match), shared only one identity (Excluded, Single Match), or did not share those identities with the featured counter-stereotypical scientist (Excluded, No Match).

These categories allowed us to test how sharing counter-stereotypical identities with featured scientists impacts the extent to which students relate to scientists. Our linear mixed model included treatment, demographic matching categories, and the interaction between the two as fixed effects. We fit our models with the R package *lme4* and specified a binomial distribution (*30*). A Type III Wald Chi-Square Test revealed significant main effects of both treatment (*X*^*2*^_2, 6244_ = 29.387, P < 0.0001) and demographic matching categories (*X*^*2*^_3, 6244_ = 15.445, P = 0.0015), as well as a significant interaction between treatment and demographic matching categories (*X*^*2*^_6, 6244_ = 15.024, P = 0.020). To directly compare students with different degrees of shared identities, we performed post-hoc Tukey pairwise comparisons and calculated Cohen’s *d* effect sizes with the R package *emmeans* (*31*). As only a subset of students provided complete demographic information, our models included 6,244 student responses from 3,102 students. We also ran a sensitivity analysis that excluded data from activities that featured white women scientists (SI Appendix).

Finally, we employed mixed effects logistic regression models to explore the effect of student-scientist demographic matching on how students responded to the open-ended prompt, “Describe how you related to the featured scientist in the activity, if at all”. For this analysis, we only included students in the humanizing treatment, as our goal was to understand how students relate to the featured scientists when the activity included humanizing information about the scientists.

For this qualitative data analysis, we first described the types of student responses to the open-ended prompt through inductive coding. We generated codes inductively from a close reading of student responses rather than searching the text of the responses for a predetermined list of categories (*32*). To create codes, two researchers independently reviewed the first term of student responses, independently developed categories that characterized student responses (i.e., codes), met to compare and revise the codes, and developed a unified codebook. Our final codebook included 24 different codes (SI Appendix, Fig. S2). After the creation of the codebook, the same two researchers used axial coding (*32*) to group and abstract codes into four categories (SI Appendix, Fig. S3). Researchers used this codebook to first independently code student responses and then collaboratively reach consensus (SI Appendix).

We integrated these codes into mixed effects logistic regression models to determine whether student-scientist demographic matching affected how the student related to the featured scientists. Specifically, we employed four different models, one for each category of student response, to measure the effect of demographic matching on the likelihood that the student 1) related to the excluded identity(s) held by the feature scientists, 2) related to the humanizing information provided in the scientist profiles, 3) related to the research interests of the featured scientists, and 4) did not relate to the featured scientists. We fit our models with the R package *lme4* and specified a binomial distribution (*30*). We included course, student ID nested with course, and quantitative biology activity as random effects in our logistic regression models. Our models included 1,324 responses from 742 students (SI Appendix).

## Data Availability

Curricular materials used in this study are available at https://datanuggets.org/dataversify/.

## Institutional Review Board and Informed Consent

This work was approved by Auburn’s Institutional Review Board (IRB) #20-603 EX 2012. We obtained informed consent from students after the nature and possible consequences of the study were explained.

## Acknowledgments

The authors would like to thank Paula Adams, Anurag Agrawal, Maria De Jesus, Ryan Dunk, Michael Smith, Jay Rosenheim, and Carl Wieman for valuable feedback. This work was funded by National Science Foundation Improving Undergraduate STEM Education grants DUE-2012014 (MGW, EHS, MKK) and DUE-2011995 (CJB, ATZ).

## Notes

### Competing Interest Statement

The authors have declared no competing interest.

