## Supporting Information for "More than a token photo: humanizing scientists enhances student engagement"

**Material Development.** We developed classroom materials that pair profiles of counter-stereotypical scientists with quantitative biology exercises (referred to as DataVersify activities hereafter) for use in this study. DataVersify activities were created by combining elements of two pre-developed science classroom materials: Data Nuggets activities (quantitative biology activities that use authentic data from real researchers in K-16 classrooms, co-designed by authors MKK and EHS, <https://datanuggets.org/>) and Project Biodiversify Scientist Profiles (researcher profiles of counter-stereotypical scientists for easy use in classrooms, co-designed by authors ATZ and MGW, www.projectbiodiversify.org). All materials used in this study are freely available for download at <https://datanuggets.org/dataversify/>.

Each DataVersify activity included three elements when presented in full: 1) a brief overview of a scientific study with associated quantitative biology questions, 2) photos of the scientist who conducted the study, and 3) humanizing information about the scientist. The Data Nugget provides real data and images from a scientific study alongside background information on the research and quantitative questions about the study for students to answer. Each Data Nugget was written in collaboration with the featured scientist. Project Biodiversify scientist profiles include both visual and humanizing elements about the scientist. Visual elements consist of two photos of the scientist - one at work and one from life outside of science, both chosen by the scientist. Examples of photos included the scientist spending time with their family, enjoying nature, and participating in Comic-Con. Project Biodiversify profiles also allow scientists to share humanizing information revealed through open-ended response prompts answered via google forms by the scientist about their life and science experiences. In the Project Biodiversify profiles, scientists responded to the following questions:

1. Why did you become a biologist?
2. What is your favorite part about your job?
3. What obstacles have you overcome to get where you are?
4. What advice do you have for aspiring biologists?
5. Do you feel that any dimension of your identity is invisible or under-represented/marginalized in STEM?
6. Can you elaborate on your answer above?

**Treatments.** Our treatments manipulated which elements of DataVersify activities were presented in classrooms across three academic terms. This approach allowed us to use the same core content and quantitative biology instruction, while varying the extent to which we shared scientist visual and humanizing content (Fig. 1A). The control treatment exposed students to Data Nuggets, but photos and Project Biodiversify scientist profiles were not included. In the visual treatment, students were exposed to Data Nuggets as well as one image of the scientist. In the humanizing treatment, students were exposed to Data Nuggets and the scientist profiles from Project Biodiversify (i.e., multiple scientist photos and humanizing information).

**Instructor Recruitment.** We recruited 43 instructors from 36 U.S. academic institutions, selected from 289 applicants after review of their course schedules and adherence to the requirements of our study design. 126 instructors met the parameters of our study, and 43 instructors accepted our invitation to participate in the study. During the study, instructors implemented materials in 37 introductory level biology courses and four upper-level courses. To recruit instructors, we used Data Nuggets and Project Biodiversify mailing lists and social media accounts, suggestions from members of our network, academic and conference listservs, and the NSF Research Coordination Network project, Equity and Diversity in Undergraduate STEM (NSF-RCN-UBE-1919462, 09/01/19-08/31/23). Our recruitment efforts resulted in a broad population of instructors teaching at R1 universities, liberal arts universities, and community colleges from 20 states (Table S5). Of the 32 instructors who provided demographic identities, 27 were women and 5 were men. This subset also included 29 white/European American, 3 Asian/Asian American, 2 Black/African American, and 1 Latino/a/x/Hispanic American instructors. All instructors were compensated for their participation in the study.

**Content Matching and Featured Counter-Stereotypical Scientist Selection.** We worked directly with each instructor to select three out of 12 DataVersify activities that best matched their course content, while still highlighting a diversity of counter-stereotypical scientists. Our aim was to center scientists with excluded identities, and, for this reason, we do not have a treatment that features white cis-man scientists, as they are not an excluded demographic group in STEM. To ensure each student in our study saw similar levels of representation, we required that each instructor included (i) a scientist who identifies as a minoritized gender (e.g., a woman or person under the trans umbrella) and (ii) a scientist of color. We chose these two categories because they are both considered to be excluded groups in STEM and, in contrast to concealable stigmatized identities, can often be, but not always, visually identified (important for the visual treatment).

**Student Population.** Our study included 3,788 students from undergraduate biology courses for majors and non-majors in the United States. We surveyed students at the beginning and end of the academic term to collect demographic information, including information about race/ethnicity and gender (Table S6). Categories for race/ethnicity are based on standards for the classification of federal data on race/ethnicity, commonly used for federal data collection purposes including the decennial census. A subset of students provided their complete demographic information (N = 3,102 students). In this subset, our student population included 1,829 white, 334 Asian, 325 Black, 299 multi-racial, 286 Latino/a/x, 20 American Indian or Alaska Native, and 9 Native Hawaiian or other Pacific Islander students. 2,002 students were women, 978 men, and 122 gender diverse. Student demographics did not differ across the three treatments (Fig. S1). The number of students who responded to our survey across all 43 instructors ranged from 11 to 246, with an average of 88 students per instructor.

**Confirmatory Factor Analysis.** Immediately after completing each DataVersify activity, we surveyed students to measure how students related to the featured scientists and engaged with the activities. Our measure of scientist relatability included one item, and our measure of student engagement included seven items, incorporating items for affective, cognitive, and behavioral engagement (Table S7) (*1*). For our measure of engagement, we specified a three-factor confirmatory factor analysis using the R package *lavaan* (*2*). Confirmatory factor analyses evaluate whether survey items are more closely related to items within their intended set (or factor) rather than to items outside of their factor (*3*). Each of our three factors included items that corresponded to either the affective (i.e., interest in the activity), cognitive (i.e., perceived importance of the activity), or behavioral (i.e., effort put into the activity) dimension of student engagement, as we expected based on the original instrument. The model fit of our three-factor CFA was good according to Chi-square statistics and common fit indices (𝜒^2^ = 708.468, df = 11, P < 0.0001; CFI = 0.982; TLI = 0.965; RMSEA = 0.092; SRMR = 0.024). Given the validity of the three engagement factors in our student sample, we used the values of these three latent variables in our CFA model as our measures of affective, cognitive, and behavioral engagement.

**Mean-Variance Standardization.** To mean-variance standardize, we calculated Z-scores using the formula Z-score = (X-μ)/s, where X is either the scientist relatability score or the engagement value measured from the CFA model, μ is the mean relatability or engagement value across the entire student sample, and s is the standard deviation around the mean. Z-scores reflect how many standard deviations a student’s score is from the average of all students across all undergraduate institutions in the sample.

**Path Analyses.** Traditional path analysis using structural equation modeling (SEM) assumes normality and independence of observations. To account for repeated measures from each student over an academic term, we employed piecewise SEM of path models using the R package *piecewiseSEM* (*4, 5*). We evaluated two separate models to explain each of the components of student engagement (affective, cognitive, and behavioral engagement): the full mediation model and the partial mediation model (Fig. S4). Piecewise SEM uses Fisher’s C statistic to evaluate the effect of missing paths (in this case, the direct path between treatment and engagement in the full mediation model) on the goodness-of-fit of the model (*4*). Models fit the data when the paths missing from the model do not differ from zero, indicated by a Fisher’s C statistic P_C_ > 0.05. All of our full mediation models fit the data (affective engagement: P_C_ = 0.189; cognitive engagement: P_C_ = 0.207; behavioral engagement: P_C_ = 0.812). We performed a chi-square difference test to evaluate whether the partial mediation models provided a better fit than the full mediation models. In the case of all three components of student engagement, we found support for the partial mediation model, despite non-significant direct links between treatment and engagement across all models (affective engagement: X^2^_diff_ = 19.875, P_diff_ < 0.0001; cognitive engagement: X^2^_diff_ = 21.092, P_diff_ < 0.0001; behavioral engagement: X^2^_diff_ = 16.432, P_diff_ < 0.0001; Table S1).

**Student-Scientist Demographic Matching.** We performed demographic matching by gender and race/ethnicity between students and the featured scientists to create the four categories described in the main text: Excluded, Double Match; Excluded, Single Match; Excluded, No Match; and Non-Excluded. We focused on gender and racial/ethnic identities because these identities are often, but not always, visually discernible (the visual treatment) and are frequently mentioned in the scientist profile (the humanizing treatment) as they relate to experiences of marginalization in STEM. We used the same survey questions (Table S6) to gather information about the gender and race/ethnicities of the students and the 12 scientists featured in the DataVersify activities. We used this information to assess whether students matched the demographics of the featured scientist. For example, if the featured scientist was a woman, students identifying as women shared the same gender identity, whereas men, transgender, nonbinary, intersex, and genderqueer students did not match the scientist’s gender. However, many individuals do not identify with these categories in a singular way. Furthermore, many gender and racial/ethnic identities are concealed identities. Because identities are complex, we made a series of decisions in the process of demographically matching students and scientists that we explain below. Our decisions are guided by our research focus on students who do not identify with the ‘stereotypical scientist’ (i.e., white cis-man).

Several of our DataVersify scientists held multiple gender identities. One of our DataVersify scientists identified as transgender and nonbinary (Table S4), and they discussed in their scientist profile the serious challenges and discrimination that transgender individuals face when pursuing educational and professional goals. Here, we considered all students holding gender diverse identities (including transgender, nonbinary, intersex, and genderqueer students) to share this gender diverse identity with this scientist, though we acknowledge each of these gender diverse identities are distinct. Another DataVersify scientist identified with both woman and intersex gender identities (Table S4). This scientist, however, did not reveal their intersex identity in their scientist profile. As this gender diverse identity was kept concealed from students, we did not consider gender diverse students to share their gender identity with this scientist.

Many students held multiple racial/ethnic identities. We matched these students with the featured scientist when the scientist identified with any of the excluded racial/ethnic identities held by the student (i.e., non-white races). However, we only matched students who exclusively identified as white with scientists who also exclusively identified as white. Two of our scientists identified with multiple racial/ethnic groups (Table S4). One scientist identified as both white and Latino/a/x and discussed their experience as the child of an immigrant from South America in their scientist profile. We considered Latino/a/x students to share their racial/ethnic identity with this scientist. However, we did not consider white students to share their racial/ethnic identity with this scientist. Another scientist identified as both white and Asian but did not mention their concealed Asian identity in their scientist profile. As such, we did not consider Asian students to share their racial/ethnic identity with this scientist. None of our scientists identified as American Indian, Alaskan Native, Native Hawaiian, or Pacific Islander (Table S4). As such, students who identified with these racial/ethnic groups did not match with one of the featured scientists in terms of race/ethnicity.

**Sensitivity Analysis.** Because four of the twelve featured scientists were white women (Table S4) and 38% of the students in our sample were white women (Fig. S1), we wondered if demographic matching between white women scientists and students drove our results. As a sensitivity analyses, we removed students who worked through activities that featured white women scientists and re-ran the linear model described above. Our model included 4,476 student responses from 2,703 students. Similar to the model above, we found significant effects of treatment (*X^2^*_2,4476_ = 28.945, P < 0.0001) and the interaction between treatment and demographic matching categories (*X^2^*_6,4476_ = 17.389, P = 0.008). However, the main effect of demographic matching categories was not significant (*X^2^*_3,4476_ = 4.853, P = 0.183). As in the model including white women scientists, students sharing both gender and race/ethnicity identities related more to the scientists in the DataVersify activities when the activities included humanizing information (*double match* - *non-excluded*: contrast: 0.531, Cohen’s *d* = 0.419, P=0.005). However, students sharing only one excluded identity did not relate more to the humanized scientists than white men students (*single match* - *non-excluded*: contrast: 0.256, Cohen’s *d* = 0.202, P = 0.133). As before, students with excluded identities who were not provided any information about the featured scientist or were exposed to only a photo of the featured scientist did not relate more to the scientists than other groups of students (control: *double match* - *non-excluded*: contrast: -0.073, Cohen’s *d* = -0.058, P=1.00; *single match* - *non-excluded*: contrast: 0.028, Cohen’s *d* = -0.011, P=1.00; visual: *double match* - *non-excluded*: contrast: -0.004, Cohen’s *d* = -0.003, P=1.00; *single match* - *non-excluded*: contrast: -0.038, Cohen’s *d* = -0.030, P=1.00).

**Qualitative Data Coding.** Over the course of our research project, five different researchers used the final codebook we created to code student responses. Co-author RMY led this qualitative coding process and coded and collaboratively reached consensus for every student response, ensuring consistency in how responses were coded. Researchers assigned responses to all appropriate codes, meaning a single response could fit in multiple codes. Every student response was coded by at least two different researchers, and each coder independently coded responses in “blocks” ranging from 20 to 1073 responses. After coding a block of responses, the two researchers met to reach 100% consensus. The average initial percent agreement for the blocks was 59%.

**Qualitative Data Analysis.** Logistic regressions revealed that the extent to which students share excluded identities with the featured scientists impacts the likelihood that students related to the excluded identity(s) of the featured scientist (*X^2^*_3, 1324_ = 59.730, P < 0.0001) and the likelihood that students did not relate at all to the featured scientist (*X^2^*_3, 1324_ = 27.633, P < 0.0001; Fig. S3). However, student-scientist demographic matching did not impact the likelihood that students related to the research interests (*X^2^*_3, 1324_ = 5.208, P = 0.157) or humanizing elements (*X^2^*_3, 1324_ = 6.155, P = 0.104) of the featured scientists (Fig. S3). Students with excluded gender and race/ethnicity identities had higher odds of relating to the excluded identities held by the featured scientists compared to students with non-excluded identities and lower odds of mentioning that they did not relate at all to the featured scientists (Fig. S3; Table S3). Over half (58.2%) of responses from students with non-excluded identities mentioned not relating to the featured scientists, whereas less than half of responses from double match (31.9%), single match (39.6%), and no match (47.6%) students mentioned not relating at all to the featured scientists (Fig. S3; Table S3).

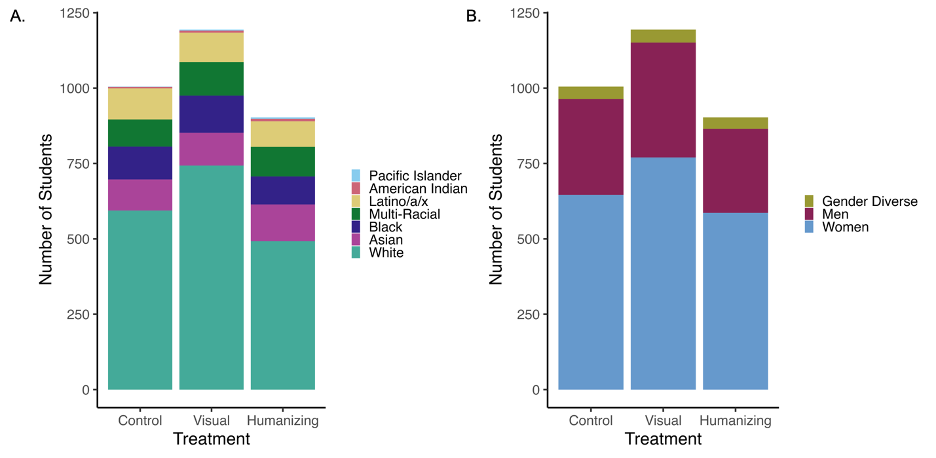

**Fig. S1. Number of students by (A) race/ethnicity and (B) gender identity across the three treatments. (A)** The control treatment was composed of 0.1% Native Hawaiian or other Pacific Islander, 0.5% American Indian or Alaska Native, 10.2% Latino/a/x, 8.9% multi-racial, 10.8% Black, 10.2% Asian, and 59.1% white students; the visual treatment was composed of 0.3% Native Hawaiian or other Pacific Islander, 0.6% American Indian or Alaska Native, 8.2% Latino/a/x, 9.3% multi-racial, 10.3% Black, 9.1% Asian, and 62.2% white students; and the humanizing treatment was composed of 0.6% Native Hawaiian or other Pacific Islander, 0.9% American Indian or Alaska Native, 9.4% Latino/a/x, 10.9% multi-racial, 10.3% Black, 13.5% Asian, and 54.5% white students. 1,829 white, 334 Asian, 325 Black, 299 multi-racial, 286 Latino/a/x, 20 American Indian or Alaska Native, and 9 Native Hawaiian or other Pacific Islander students. **(B)** The control treatment was composed of 4.1% gender diverse, 31.6% men, and 64.3% women students; the visual treatment was composed of 3.6% gender diverse, 31.9% men, and 64.5% women students; and the control treatment was composed of 4.2% gender diverse, 30.1% men, and 64.9% women students.

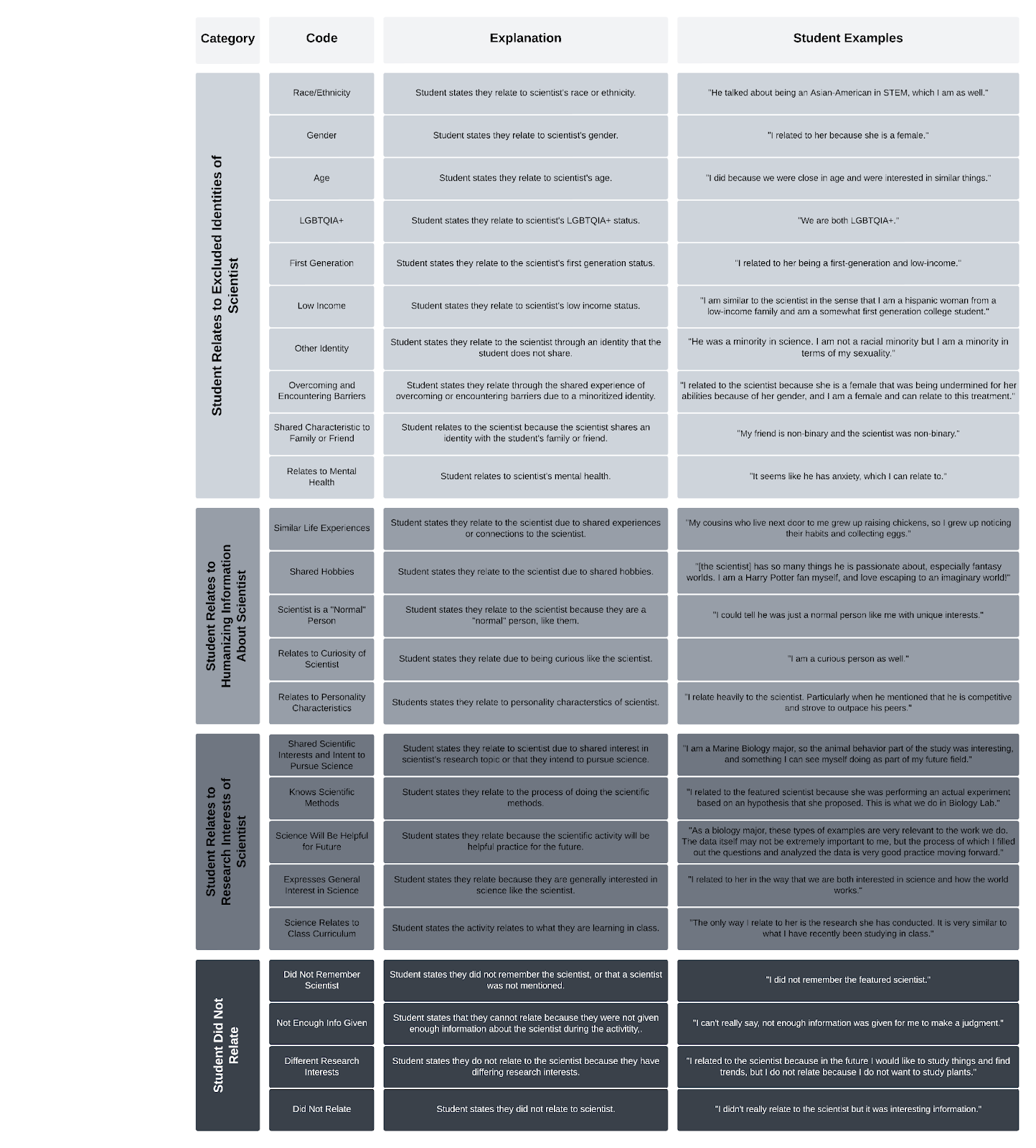

**Fig. S2. Codebook for student responses to the open-ended response “Describe how you related to the featured scientist in the activity, if at all”.** The codebook includes 24 different codes, grouped into 4 categories.

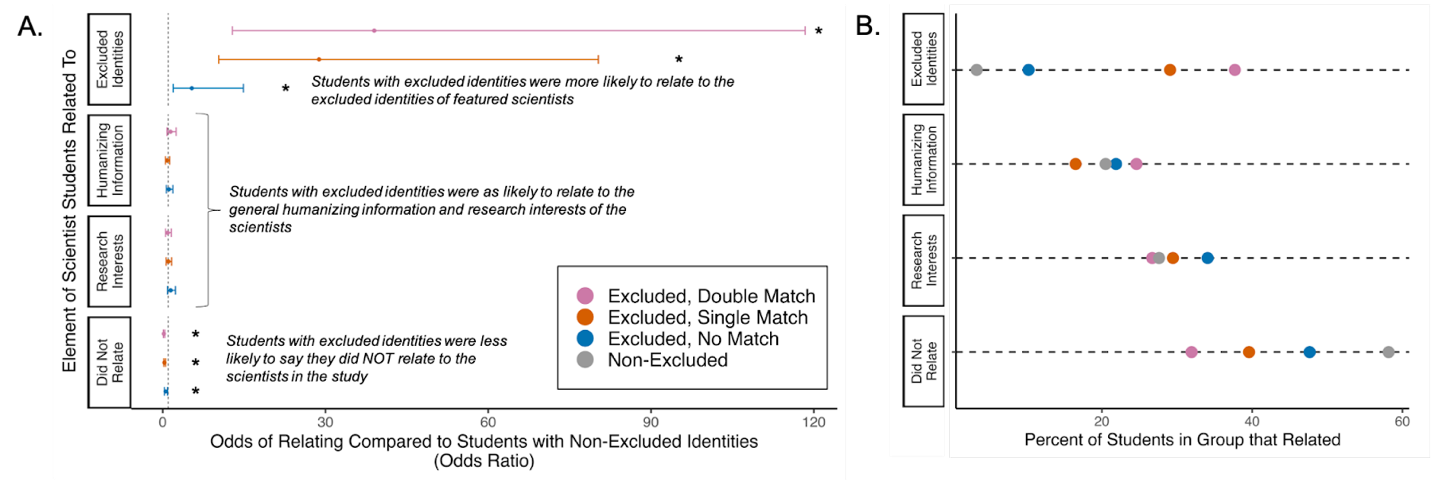

**Fig. S3. What about scientists did students with historically excluded identities relate to?** **(A)** Students with excluded identities had higher odds of relating to the excluded identities held by the featured scientists and lower odds of not relating at all to the featured scientists compared to students with non-excluded identities. Odds ratios and 95% confidence intervals of students with excluded identities across each of the four categories characterizing student responses to the open-ended prompt “Describe how you related to the featured scientist in the activity, if at all”. The reference group included non-excluded students (i.e., white men). Confidence intervals that do *not* cross the dashed line at x = 1 are statistically significant, indicated by asterisks. **(B)** The percent of students for each demographic matching category that related to aspects of the featured scientist for each of the four categories.

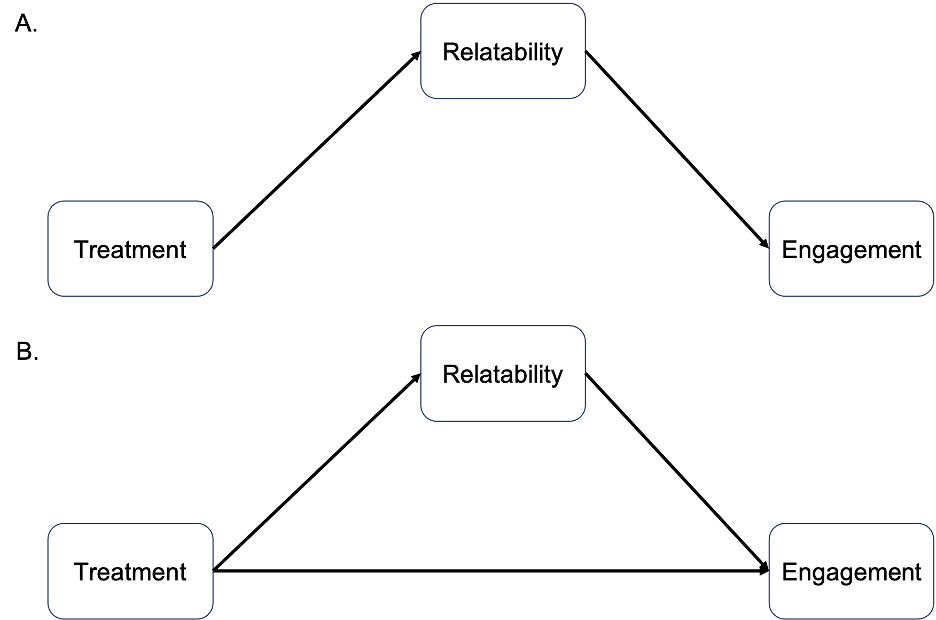

**Fig. S4. Depictions of full (A) and partial (B) mediation models.** **(A)** The full mediation model posits that scientist relatability fully mediates the relationship between treatment and student engagement. **(B)** The partial mediation model posits that a direct relationship exists between treatment and student engagement, in addition to a mediated relationship between treatment and student engagement. We ran three different full and partial mediation models, one for each component of student engagement (affective, cognitive, and behavioral engagement).

**Table S1.** Estimates, standard errors (SE), and significance values for paths included in the partial path models. Each partial path model included a different component of student engagement: affective, cognitive, and behavioral engagement. Mean-variance standardized estimates are reported for continuous variables and marginal means are reported for categorical variables. Significant P-values at the α=0.05 level are bolded.

| **Model** | **Path** | **Estimate** | **SE** | **P-value** |
| --- | --- | --- | --- | --- |
| **All** | **Treatment to Relatability** | **Control = -0.074** | **0.037** | **<0.0001** |
|  |  | **Visual = -0.040** | **0.035** |  |
|  |  | **Humanizing = 0.124** | **0.037** |  |
| **Affective Engagement** | **Relatability to Affective Engagement** | **0.370** | **0.010** | **<0.0001** |
|  | Treatment to Affective Engagement | Control = 0.016 | 0.051 | 0.370 |
|  |  | Visual = -0.019 | 0.050 |  |
|  |  | Humanizing = -0.046 | 0.051 |  |
| **Cognitive Engagement** | **Relatability to Cognitive Engagement** | **0.378** | **0.010** | **<0.0001** |
|  | Treatment to Cognitive Engagement | Control = 0.029 | 0.039 | 0.565 |
|  |  | Visual = -0.020 | 0.038 |  |
|  |  | Humanizing = -0.023 | 0.039 |  |
| **Behavioral Engagement** | **Relatability to Behavioral Engagement** | **0.289** | **0.011** | **<0.0001** |
|  | Treatment to Behavioral Engagement | Control = -0.008 | 0.042 | 0.480 |
|  |  | Visual = -0.013 | 0.041 |  |
|  |  | Humanizing = -0.030 | 0.043 |  |

**Table S2.** Summary of pairwise comparisons between all student-scientist demographic matching categories for each treatment group from the linear mixed model estimating the effect of student-scientist demographic matching on scientist relatability. Differences in estimated marginal means (contrast estimate), standard errors of the estimates (SE), Cohen’s *d* effect sizes, and significance values from post-hoc Tukey pairwise comparisons are reported. P-values were adjusted using the Tukey method for comparing a family of 12 estimates. Significant P-values at the α=0.05 level are bolded.

| **Treatment** | **Contrast** | **Contrast Estimate** | **SE** | **Effect Size (Cohen's *d*)** | **P-value** |
| --- | --- | --- | --- | --- | --- |
| **Humanizing** | Double Match - Single Match | 0.109 | 0.066 | 0.087 | 0.900 |
|  | Double Match - No Match | 0.185 | 0.069 | 0.147 | 0.245 |
|  | **Double Match - Non-Excluded** | **0.416** | **0.093** | **0.331** | **0.0005** |
|  | Single Match - No Match | 0.076 | 0.051 | 0.061 | 0.942 |
|  | **Single Match - Non-Excluded** | **0.307** | **0.080** | **0.245** | **0.007** |
|  | No Match - Non-Excluded | 0.231 | 0.083 | 0.184 | 0.189 |
| **Visual** | Double Match - Single Match | 0.067 | 0.059 | 0.053 | 0.993 |
|  | Double Match - No Match | 0.104 | 0.061 | 0.083 | 0.867 |
|  | Double Match - Non-Excluded | 0.075 | 0.078 | 0.060 | 0.999 |
|  | Single Match - No Match | 0.037 | 0.045 | 0.030 | 1.000 |
|  | Single Match - Non-Excluded | 0.008 | 0.068 | 0.006 | 1.000 |
|  | No Match - Non-Excluded | -0.029 | 0.070 | -0.023 | 1.000 |
| **Control** | Double Match - Single Match | -0.028 | 0.060 | -0.022 | 1.000 |
|  | Double Match - No Match | -0.041 | 0.066 | -0.033 | 1.000 |
|  | Double Match - Non-Excluded | 0.071 | 0.083 | 0.057 | 1.000 |
|  | Single Match - No Match | -0.013 | 0.049 | -0.010 | 1.000 |
|  | Single Match - Non-Excluded | 0.099 | 0.072 | 0.079 | 0.969 |
|  | No Match - Non-Excluded | 0.112 | 0.077 | 0.089 | 0.950 |

**Table S3.** Summary of four mixed effect logistic regression models, each exploring the effect of student-scientist demographic matching on how students responded to the open-ended prompt: “Describe how you related to the featured scientist in the activity, if at all”. Each model included a different category of student response to the prompt: students either did not relate to the featured scientist, related to the scientist’s research interests, related to humanizing information about the scientist, and/or related to the excluded identity of the scientist. Regression coefficient estimates (estimate), standard errors of the estimates (SE), odds ratios, percentages of student responses within each demographic matching category, and significance values are reported. The reference group is non-excluded students (i.e., white men). Significant P-values at the α=0.05 level are bolded.

| **Model** | **Fixed Effect** | **Estimate** | **SE** | **Odds Ratio** | **Percent** | **P-value** |
| --- | --- | --- | --- | --- | --- | --- |
| **Did Not Relate** | **Double Match** | **-1.530** | **0.326** | **0.216** | **31.9%** | **<0.0001** |
|  | **Single Match** | **-1.129** | **0.263** | **0.323** | **39.6%** | **<0.0001** |
|  | **No Match** | **-0.645** | **0.273** | **0.524** | **47.6%** | **0.018** |
| **Relate to Research Interests** | Double Match | -0.101 | 0.284 | 0.903 | 26.7% | 0.721 |
|  | Single Match | 0.032 | 0.232 | 1.033 | 29.5% | 0.889 |
|  | No Match | 0.372 | 0.244 | 1.450 | 34.1% | 0.128 |
| **Relate to Humanizing Information** | Double Match | 0.349 | 0.283 | 1.418 | 24.6% | 0.217 |
|  | Single Match | -0.217 | 0.240 | 0.805 | 16.5% | 0.366 |
|  | No Match | 0.134 | 0.249 | 1.143 | 21.9% | 0.591 |
| **Relate to Excluded Identities** | **Double Match** | **3.663** | **0.557** | **38.981** | **37.7%** | **<0.0001** |
|  | **Single Match** | **3.361** | **0.523** | **28.807** | **29.1%** | **<0.0001** |
|  | **No Match** | **1.678** | **0.522** | **5.354** | **10.2%** | **0.0013** |

**Table S4.** Summary of identities represented by the twelve scientists profiled in the DataVersify activities, and the number of times an activity with that identity was implemented in the study. Scientists completed a survey to identify their gender/sex and race/ethnicity identities and were able to select multiple identities per identity category.

| **Scientist Identity** | | **Number of Featured Scientists** | **Number of Implementations in Study** |
| --- | --- | --- | --- |
| **Gender/Sex** | Man | 3 | 34 |
|  | Woman | 8 | 81 |
|  | Transgender | 1 | 16 |
|  | Nonbinary | 1 | 16 |
|  | Intersex | 1 | 11 |
| **Race/**  **Ethnicity** | Black/African American | 2 | 19 |
|  | Asian/Asian American | 3 | 32 |
|  | Latino/Hispanic/Hispanic American | 2 | 18 |
|  | White/European American | 7 | 77 |

**Table S5.** Information about the thirty-six US undergraduate institutions that participated in our study was acquired from Carnegie Classifications ([https://carnegieclassifications.acenet.edu/)](https://carnegieclassifications.acenet.edu/). We used Wikipedia ([www.wikipedia.org](http://www.wikipedia.org)) when information was not available from Carnegie Classifications.

| **Institution Type** | Private | 2 |
| --- | --- | --- |
|  | Private Not-For-Profit | 7 |
|  | Public | 27 |
| **Institution Year** | 2-Year | 10 |
|  | 4-Year | 26 |
| **Institution Regional Location** | Northeast | 6 |
|  | Midwest | 7 |
|  | South | 15 |
|  | West | 8 |

**Table S6.** Survey questions used to collect the race/ethnicities and genders of the students participating in this study and the scientists featured in the DataVersify activities. We defined students and scientists as having excluded identities if they identified as American Indian/Alaska Native, Asian, Black, Latino, Native Hawaiian or other Pacific Islander, women, intersex, transgender, agender, nonbinary, and/or genderqueer individuals. Students filled out this demographic information at the beginning and end of the course and responded to questions about their gender before questions about their race.

**What term best describes your gender identity? (select all that apply)**

Man

Woman

Intersex

Transgender

Nonbinary

Genderqueer

My gender is:

**What is your race*? (select all that apply)**

American Indian/Alaska Native

Asian/Asian American

Black/African American

Latino/Hispanic/Hispanic American

Native Hawaiian or Other Pacific Islander

White/European American

My race is:

*According to the National Institutes of Health (NIH), we define each racial category as follows:

**American Indian or Alaska Native.** A person having origins in any of the original peoples of North and South America (including Central America), and who maintains tribal affiliation or community attachment.

**Asian/Asian American.** A person having origins in any of the original peoples of the Far East, Southeast Asia, or the Indian subcontinent including, for example, Cambodia, China, India, Japan, Korea, Malaysia, Pakistan, the Philippine Islands, Thailand, and Vietnam.

**Black/African American.** A person having origins in any of the black racial groups of Africa.

**Latino/Hispanic/Hispanic American.** A person of Cuban, Mexican, Puerto Rican, South or Central American, or other Spanish culture or origin, regardless of race. The term, “Spanish origin,” can be used in addition to “Hispanic or Latino.”

**Native Hawaiian or Other Pacific Islander.** A person having origins to the original peoples of Hawaii, Guam, Samoa, or other Pacific Islands.

**White/European American.** A person having origins in any of the original people of Europe, the Middle East, or North Africa.

**Table S7.** Survey items on a 7-point sliding scale were developed using the Experience Sampling Method (*8*) to measure student engagement with the activity and student relatability to the featured scientist. Students completed surveys immediately after completing the activity and responded to items about their engagement with the activity before items about how they related to the featured scientist.

**Engagement with Activity (*9*)**

***Behavioral Engagement***

How hard were you working on the activity?

How well were you concentrating on the activity?

***Cognitive Engagement***

How important was the activity to you?

How important was this activity to your future?

***Affective Engagement***

Was the activity interesting?

Did you enjoy the activity?

How engaging did you find the activity?

**Relatability of Featured Scientist**

If a scientist was mentioned, to what extent did you relate to the featured scientist in the activity?
